# A role for DNA methylation in bumblebee morphogenesis hints at female-specific developmental erasure

**DOI:** 10.1101/2023.05.26.542467

**Authors:** B. J. Hunt, M. Pegoraro, H. Marshall, E. B. Mallon

## Abstract

Epigenetic mechanisms, such as DNA methylation, are crucial factors in animal development. In some mammals almost all DNA methylation is erased during embryo development and re-established in a sex- and cell-specific manner. This erasure and re-establishment is thought to primarily be a vertebrate-specific trait. Insects are particularly interesting in terms of development as many species often undergo remarkable morphological changes en route to maturity, i.e. morphogenesis. However, little is known about the role of epigenetic mechanisms in this process across species. We have used whole genome bisulfite sequencing to track genome-wide DNA methylation changes through the development of an economically and environmentally important pollinator species, the bumblebee Bombus terrestris. We find overall levels of DNA methylation vary throughout development and we find developmentally relevant differentially methylated genes throughout. Intriguingly, we have identified a depletion of DNA methylation in ovaries and an enrichment in sperm. We suggest this could represent a sex-specific DNA methylation erasure event. To our knowledge this is the first suggestion of possible developmental DNA methylation erasure in an insect species. This study lays the required groundwork for functional experimental work to determine if there is a causal-nature to the DNA methylation differences identified. Additionally, the application of single-cell methylation sequencing to this system will enable more accurate identification of when/if DNA methylation is erased during development.

## Introduction

Epigenetic mechanisms such as DNA methylation, chromatin modifications and non-coding RNA silencing can cause stable and heritable changes in gene expression. They are also crucial factors in animal development, specifically in generating cell-specific gene expression patterns from a single genome (Paulsen and Ferguson-Smith, 2001). The most well-researched of these mechanisms is DNA methylation, which most commonly involves the addition of a methyl group to a cytosine nucleotide in animals, primarily in a CpG context. In mammals almost all DNA methylation is erased during embryo development and re-established in a sex- and cell-specific manner (Seisenberger *et al*., 2012), aside from any DNA methylation mediating imprinted genes, and this re-programming is thought to primarily be a vertebrate-specific trait (Xu *et al*., 2019). While this process is well-understood in mammalian model lab species, we know little about how DNA methylation functions in the development of other clades. Insects are particularly interesting in terms of development as many species often undergo remarkable morphological changes en route to maturity.

The function of DNA methylation across insects is highly variable (Lewis *et al*., 2020), with high levels of gene-body DNA methylation generally linked to the stable expression of housekeeping genes (Provataris *et al*., 2018). Insects show significantly different DNA methylation profiles compared to mammals, with considerably lower genome-wide levels of DNA methylation, from less than one percent to around fourteen percent depending on the species (Bewick *et al*., 2016), and generally exhibiting only gene-body DNA methylation (Provataris *et al*., 2018), although see Lewis *et al*. (2020). Within insects, there are also notable differences in DNA methylation levels between holometabolous species and hemimetabolous species, with holometabolous insects generally showing lower levels of DNA methylation in the adult stage (Bewick *et al*., 2016).

Holometabolous undergo a process called metamorphosis during development, defined as an extreme change in body form/structure throughout development. Holometabolous insects such as butterflies, beetles and bees, develop from eggs into larvae, which transform into pupae before reaching a phenotypically different adult stage. Unlike the well characterised epigenetic dynamics of mammalian development, the processes underpinning insect development remain more elusive. Currently only a handful of studies have examined DNA methylation dynamics throughout insect development. In the red flour beetle, *Tribolium castaneum*, and the tobacco hornworm, *Manduca sexta*, DNA methylation has been show to decrease during embryo development with levels increasing in the final adult stage (Song *et al*., 2017; Gegner *et al*., 2021). *Drosophilia melanogaster* displays little to no DNA methylation in the adult stage and lacks the essential genes involved in DNA methylation establishment and maintenance (DNMT1 and DNMT3), however DNA methylation was found to be present in slightly elevated levels in the embryo and pupal stages of development (Deshmukh *et al*., 2018). The primary focus of developmental epigenetics within insects, however, is within the eusocial species.

Social insects display extreme phenotypic differences between castes, e.g. workers/queens/drones, even though individuals share highly similar and sometimes identical genetic backgrounds. Epigenetic processes are therefore thought to be fundamental to many of the phenotypic differences observed. It was recently noted that there is conflicting evidence for a functional role of DNA methylation in caste determination in social insects (Oldroyd and Yagound, 2021), although many of these studies focus solely on final adult stages. Harris *et al*. (2019) have recently explored the DNA methylome throughout honeybee development, finding fluctuating levels of DNA methylation between developmental stages, with larvae showing the lowest DNA methylation levels and sperm showing the highest. Bonasio *et al*. (2012) explored developmental DNA methylation in two ant species, finding the lowest DNA methylation levels were in the embryo of one species (*Camponotus floridanus*) and the larvae of another (*Harpegnathos saltator*).

Therefore evidence exists for a role of DNA methylation during metamorphosis in insects, and specifically within social insects, but the extent to which this is conserved between related species remains unknown. Interestingly, even within mammals the dynamic re-programming of DNA methylation marks during development differs between species (Beaujean *et al*., 2004). To better understand the mechanisms driving development and metamorphosis in insects we must explore these processes in a variety of independent species. Additionally, DNA methylation in invertebrates has recently attracted attention as a possible mechanism of environmental adaptation. Determining the extent of DNA methylation developmental re-programming in insects will have profound implications for the role of environmentally induced DNA methylation heritability, and thus the field of adaptive epigenetics.

Bumblebees provide an ideal model to explore the role of DNA methylation during development as they possess a functional DNA methylation system (Sadd *et al*., 2015) which may play a regulatory role in social caste determination (Amarasinghe *et al*., 2014; Li *et al*., 2018; Marshall *et al*., 2019). They are also both economically and environmentally important as a pollinator species (Woodard *et al*., 2015). *Bombus terrestris* colonies are founded annually by a singly mated queen; she will initially lay diploid eggs resulting in female workers (Bloch, 1999). A competition phase later occurs between queens and workers, where some workers will become reproductive, without mating, and produce their own haploid sons (Alaux *et al*., 2006). We take advantage of this system to follow whole-genome and gene-level DNA methylation dynamics through female worker somatic tissue (brain), ovary tissue, male offspring larval head tissue, male pupal head tissue, adult male brain tissue and finally sperm. If DNA methylation does play a role in the metamorphosis of *B. terrestris* we would predict; (i) global differences in DNA methylation levels between different developmental stages and (ii) differentially methylated genes involved in developmental processes between developmental stages.

## Methods

### Husbandry and tissue dissection

Five colonies of *Bombus terrestris audax* (Agralan, UK) were established and maintained in wooden nest boxes under red light at at 26°C and 60% humidity on a diet of 50% v/v glucose/fructose apiary solution (Meliose-Roquette, France) and pollen (Percie du set, France) (Amarasinghe *et al*., 2014). Between 80 - 100 days after establishment, callow females were isolated in perspex boxes (18.5 cm x 12.5cm x 6.5cm) for three days, at which point a further two callow females were added, with old and new callows marked in different colours to identify them. When the older female assumed a dominant role and began laying eggs, larvae and pupae samples were collected (Figure 1). The reproductive female’s brain and ovaries were also sampled. For male brain and sperm samples, offspring males were collected as callows and kept in groups of 10 in a perspex box for 13 - 16 days before dissection and extraction (Baer and Schmid-Hempel 2000). All samples were collected in a 2 hour window between 3 - 5pm to avoid circadian influences on the methylome.

**Figure 1.**
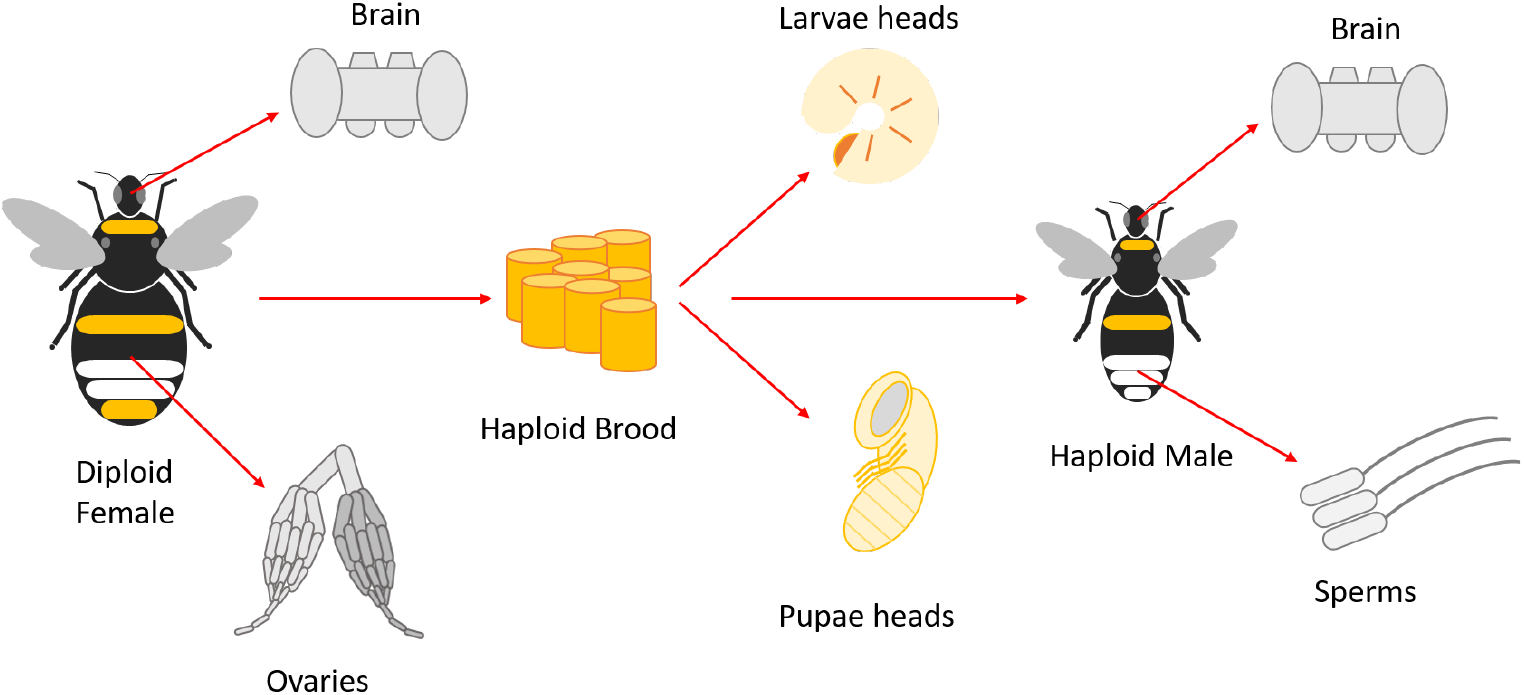
Schematic overview of the samples collected. Specifically, brain and ovary tissue were taken from female workers. Larval heads, pupal heads, brain tissue and sperm were then taken from their male offspring.

Reproductive workers and adult males were anaesthetised on ice (4°C) and brains/ovaries dissected in PBS. For sperm collection the protocol described in Baer and Schmid-Hempel (2000) was followed, collecting sperm in 20*µ*l PBS from accessory testis excluding the testis tissue. Heads were dissected from larvae stage four and from pupae with eye pigmentation but no body pigmentation. Samples were snap-frozen in liquid nitrogen and stored at -80°C.

### Genomic DNA extraction, sequencing and mapping

Twenty libraries were prepared from 3 colonies, comprising four replicates each of reproductive workers brains, ovarian tissue, larval heads and pupal heads, and two replicates each of male brains and sperm. Genomic DNA was extracted using QIamp DNA Micro Kit (QIAGEN) with minor modifications. Prior to the addition of Buffer ATL and Proteinase K, tissues were crushed in liquid nitrogen-chilled eppendorf tubes using a mini-pestle. Carrier RNA was used with Buffer AL. DNA concentration was assessed using Nanodrop and Qubit® dsDNA BR Assay Kit (ThermoFisher Scientific, USA). Whole genome bisulfite sequencing (WGBS), i.e. library preparation, bisulfite conversion and 150bp paired-end sequencing using a HiSeq 2000 machine (Illumina, Inc.) was performed at the Beijing Genomics Institute (BGI). All samples include a 1% lambda spike to assess bisulfite conversion efficiency.

Reads were subject to quality control using Trim Galore 0.6.5 (clip_R1 10 –clip_R2 10). Bismark 0.20.0 (Krueger and Andrews, 2011) was used to align the reads to the Bter_1.0 genome (Refseq accession no. GCF_000214255.1 (Sadd *et al*., 2015)) (–score-min L,0,-0.4) and remove PCR artifacts in the CpG context. To address an issue with unreliable calls at certain base positions due to flowcell tile quality, deduplicated BAM files were processed using the methylKit 1.16.1 (Akalin *et al*., 2012) function processBismarkAln to exclude bases with minqual < 30. The output of this was reformatted to the Bismark coverage format using R 4.0.3 in RStudio 1.3.1093, and then fed back into Bismark’s coverage2cystosine module using the option merge_CpGs to destrand the data and generate a whole genome CpG methylation call file per sample.

The sequenced samples returned libraries ranging from 24 - 38 million paired end reads with alignment rates from 53 - 80%. After deduplication this gave an average genome-wide coverage of 14X ±3X. The mean number of methylated cytosines in a CpG context was 0.71% ± 0.07% (standard deviation), in a CHG context 0.45% ± 0.03% and in a CHH context 0.47% ± 0.03%. Methylation levels in the latter two contexts did not exceed the error rate (0.5%) estimated by alignment of the libraries to the lambda spike-in genome. These levels of DNA methylation are in-line with previous work in this species (Marshall *et al*., 2019).

### DNA methylation differences between developmental stages

Differential methylation analysis was performed using methylKit (Akalin *et al*., 2012). The destranded calls were imported using the mincov parameter to exclude sites with coverage < 10 reads, and further filtered to exclude sites with coverage higher than the 99.5th percentile. Differential methylation analyses were conducted between worker brain - worker ovaries; worker ovaries - larvae head; larvae head - pupae head; pupae head - male brain; male brain - sperm; worker brain - male brain; worker ovaries - sperm. For each comparison, methylKit’s reorg function was used to extract the relevant samples and reassign treatment IDs. In comparisons where four replicates per group were available for both groups, sites were required to be represented in at least three replicates per group to be tested. In comparisons where two replicates were available in one or both groups, sites were required to be represented in two replicates per group to be tested. A binomial test was used to make per-loci methylation status calls using a 0.5% error rate estimated from the sequencing lambda spike-in control and only loci identified as methylated in at least one sample were tested for differential methylation using the Chi-squared test, controlling for colony as a covariate and correcting for overdispersion. Loci were considered to be differentially methylated if they show a methylation difference of 10% or greater and an FDR-adjusted q-value of 0.05. Loci were annotated with bedtools intersect (Quinlan and Hall, 2010). To then call a gene differentially methylated, we required an exon within a given gene to contain at least one significant differentially methylated CpG in addition to having an overall methylation difference exon-wide of 15%.

Genome-wide DNA methylation profiles were calculated as the mean weighted methylation level (Schultz *et al*., 2012) across samples per developmental stage for each genomic feature. Differences in overall gene-level DNA methylation were determined using linear mixed effects models implemented from the lme4 R package 1.1-33 where replicate and gene were random factors. Post-hoc testing was done with the emmeans R package 1.8.6. Genomic features were classed as showing high methylation (weighted methylation >0.7), medium methylation (>0.3-0.7), low methylation (>0-0.3) or no methylation. Introns, 5’ UTR and 3’ UTR regions were annotated using AGAT 0.8.0 (Dainat, 2021). Putative promoter regions were classed as 500bp upstream of each gene’s 5’ UTR and intergenic regions were all other unannotated regions, although we acknowledge these regions likely include transposable elements. The extent of variability in DNA methylation levels across developmental stages was assessed using the Jensen-Shannon divergence (JSD) index implemented by the MethylIT 0.3.2.3 R package (Sanchez, 2021). Genes were classed as showing significant variability across developmental states when the JSD index was >0.037 based on the distribution of all indices (supplementary Fig.S1).

### GO term enrichment

Gene ontology (GO) terms for *B. terrestris* were taken from a custom database made in Bebane *et al*. (2019). GO enrichment analysis was carried out using the hypergeometric test with Benjamini-Hochberg (Benjamini and Hochberg, 1995) multiple-testing correction, q <0.05. GO terms from differentially methylated genes between comparisons were tested against a GO term database made from the GO terms associated with all methylated genes per comparison. Genes were determined as methylated if they had a mean weighted methylation level greater than the bisulfite conversion error rate of 0.05 in any of the samples within each comparison. Genes classed as highly methylated per developmental stage were tested for enrichment against all methylated genes for that developmental stage. Genes classed as showing no methylation per developmental stage were tested against all genes present in the WGBS for that developmental stage. Genes with a significant JSD index were tested against all genes present across all data sets which were methylated in at least one developmental stage. REVIGO (Supek *et al*., 2011) was used to generate GO descriptions from the GO ids.

## Results

### Genome-wide developmental DNA methylation dynamics

Genome-wide DNA methylation patterns are most different in worker ovaries and sperm compared to brain/head tissue of adults and developing larvae/pupae (Fig. 2a). As DNA methylation in enriched in gene-bodies in bumblebees we assessed global gene levels across developmental stages. Using a linear mixed effects model we find all developmental stages show significant differences in gene-level DNA methylation, apart from reproductive worker brains and larvae heads (Fig. 2b, supplementary 1.0.0). Specifically, sperm shows the highest levels of DNA methylation in genes and female ovaries the lowest (Fig. 2b, supplementary 1.0.0).

**Figure 2.**
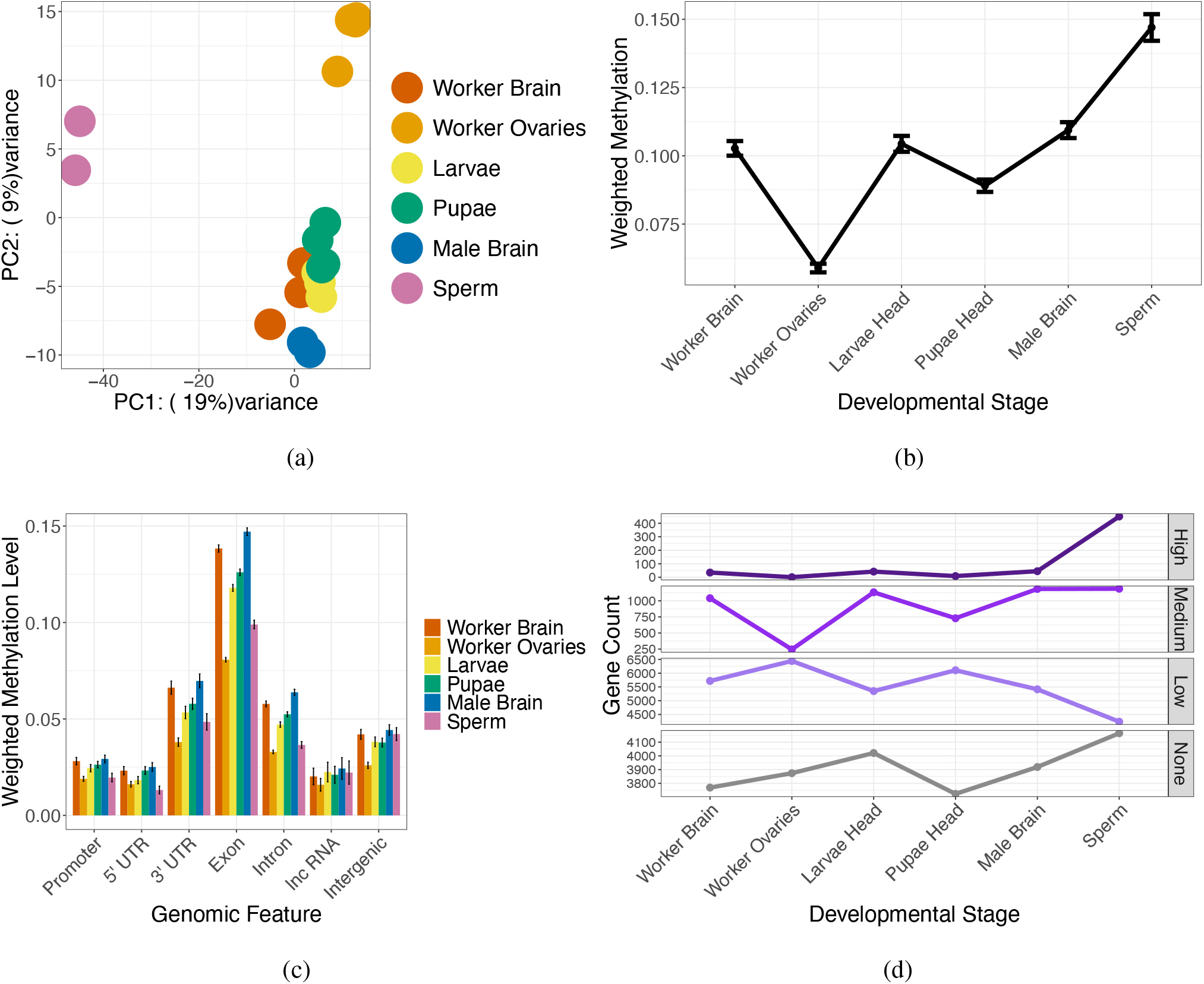
(a) PCA plot of the methylation level of CpGs which have data in all samples and are classed as methylated in at least one sample via binomial test (n = 1,189). (b) Gene levels of DNA methylation across developmental stages (n = 8,874 - 10,549). Each dot represents the mean methylation level across all genes and replicates per developmental stage. The error bars represent 95% confidence intervals. (c) Mean level of DNA methylation per genomic feature averaged across all replicates per developmental stage. The error bars represent 95% confidence intervals. (d) Number of genes per developmental stage that are classed as showing high methylation (>0.7), medium methylation (>0.3-0.7), low methylation (>0-0.3) and no methylation.

To further explore this pattern genome-wide we also looked at DNA methylation levels across different genomic features for each developmental stage. We find across stages that DNA methylation is enriched in exons and depleted in promoter and 5’ UTR regions, in line with previous findings in adult somatic tissue (Lewis *et al*., 2020). The depletion of DNA methylation in worker ovaries observed in Figure 2b is present genome-wide across all genomic features (supplementary Fig. S2).

We also find that the high mean level of DNA methylation in sperm genes is driven by sperm containing substantially more highly methylated genes compared to all other developmental stages (Fig.2d). Interestingly, sperm also contains more genes with no DNA methylation compared to all other developmental stages, suggesting a more bimodal pattern of DNA methylation where DNA methylation is specifically targeting or being erased from certain genes (Fig.2d). In order to explore the function of highly methylated genes and genes with no DNA methylation in all developmental stages we carried out a GO enrichment analysis. Highly methylated genes in all developmental stages are enriched for a variety of core cellular processes (supplementary 1.0.1, 1.0.2), consistent with a role for DNA methylation in maintaining housekeeping gene function, whereas genes with no DNA methylation across developmental stages are involved in a much larger variety of metabolic and cellular processes (supplementary 1.0.1, 1.0.3).

### Differential DNA methylation between developmental stages

In order to examine in which genes DNA methylation changes throughout bumblebee development, we first identified significanly differentially methylated CpGs per sequential comparison. We find the vast majority of significantly differentially methylated CpGs in all comparisons are found within exon regions (supplementary Fig. S3 and S4) and are equally distributed across exons within a gene (supplementary Fig. S5). The number of hypermethylated (i.e. having more methylation) CpGs per comparison mirrors the global DNA methylation trends between developmental stages as seen in Fig. 2b, e.g. there are significantly more hypermethylated CpGs in worker brains compared to worker ovaries (supplementary table S1).

To class a gene as differentially methylated between developmental stages we required the gene to contain an exon which possessed both a significantly differentially methylated CpG and an overall weighted methylation difference of 0.15 (equivalent to 15% overall methylation difference) in the same direction as the CpG, i.e. if the CpG is hypermethylated in worker brains compared to ovaries the entire exon must also be hypermethylated in worker brains compared to ovaries. Again we find the number of differentially methylated genes mirrors the overall trends in DNA methylation seen in Figure 2b (Table 1). We also checked the overall levels of methylation for genes which are classed as differentially methylated. We find the majority of differentially methylated genes show overall low levels of DNA methylation, i.e. have a weighted methylated level greater than 0 and less than or equal to 0.3 (supplementary Fig.S6).

**Table 1:** The number of genes with a hypermethylated exon per developmental stage for all comparisons. A chi-squared goodness of fit was carried out per comparison to determine if there are significantly more hypermethylated genes in one developmental stage compared to the other.

|  | **Worker Brain** | **Worker Ovaries** | **Larvae Head** | **Pupae Head** | **Male Brain** | **Sperm** | **X-squared** | **df** | **p** |
| --- | --- | --- | --- | --- | --- | --- | --- | --- | --- |
| **Worker Brain vs Worker Ovaries** | 1220 | 2 |  |  |  |  | 1214 | 1 | <0.001* |
| **Worker Ovaries vs Larvae Head** |  | 1 | 285 |  |  |  | 282.01 | 1 | <0.001* |
| **Larvae Head vs Pupae Head** |  |  | 4 | 1 |  |  | 1.8 | 1 | 0.18 |
| **Pupae Head vs Male Brain** |  |  |  | 4 | 153 |  | 141.41 | 1 | <0.001* |
| **Male Brain vs Sperm** |  |  |  |  | 2 | 47 | 41.32 | 1 | <0.001* |
| **Worker Brain vs Male Brain** | 12 |  |  |  | 25 |  | 4.56 | 1 | 0.032* |
| **Worker Ovaries vs Sperm** |  | 0 |  |  |  | 134 | 134 | 1 | <0.001* |

Across all comparisons, there are a total of 1,516 genes which change methylation level during development (supplementary 1.0.4). The majority of these (962) are unique to the comparison between worker brains and ovaries (supplementary Fig. S9). A GO enrichment analysis was then carried out for differentially methylated genes in each comparison, compared against a background set of all methylated genes found within those specific samples. In all comparisons, we find a wide variety of GO terms enriched between developmental stages (supplementary 1.0.5). Of particular note, we specifically find GO terms involved in neuron development and oogenesis enriched between worker brain and ovary tissue. Between male and worker female brains we find the GO terms “*sex di*.*fferentiation*” (GO:0007548), “*gonad development*” (GO:0008406) and “*development of primary male sexual characteristics*” (GO:0046546) enriched. We also find the terms “*instar larval or pupal development*” (GO:0002165) and “*imaginal disc pattern formation*” (GO:0007447) enriched between ovaries and larvae. Between ovaries and sperm we find “*female gamete generation*” (GO:0007292) and “*cell fate determination*” (GO:0001709) enriched. Finally, differentially methylated genes throughout developmental comparisons contain GO terms enriched for alternative splicing. See supplementary 1.0.5 for all enriched GO terms.

### Variability of DNA methylation across development

In order to identify genes with the most variable DNA methylation levels throughout development, we calculated the Jensen-Shannon Diversity index per gene per comparison and took the mean index across all comparisons. We find 827 genes show a considerably higher than average variability in methylation levels (supplementary 1.0.6, Fig.S1). These genes are distributed evenly across the genome (supplementary Fig.S7), with a slightly higher proportion of genes showing a significant JSD index on chromosome NC_015774.1 (supplementary Fig.S7).

In order to explore the function of these genes, we carried out a GO enrichment analysis of the significantly variable genes compared to all genes present in the data set. We find mostly metabolic related processes enriched, however there are also some terms enriched related to RNA splicing (supplementary 1.0.6). We then looked for GO enrichment in the significantly variable genes on chromosome NC_015774.1 compared to all significantly variable genes finding various processes including “*sperm individualization*” (GO:0007291) and “*determination of adult lifespan*” (GO:0008340), supplementary 1.0.6. We find two genes specifically driving the enrichment of the lifespan-related GO term. The first is *ras-like protein 1* (LOC100644593), which is involved in cell-division and growth and is required for normal development (Bollag and McCormick, 1997). This gene is more highly methylated in the larval head tissue and sperm (supplementary Fig. S8). The second gene, *caspase-6* (LOC100652135) controls cell-death and the inflammation response (Cohen, 1997; Bartel *et al*., 2017). This gene is more highly methylated in head and brain tissue throughout development, compared to gametes (supplementary Fig. S8).

We checked if the highly variable genes are also significantly differentially methylated between stages. We find 24% of variable genes are also differentially methylated (n = 200), compared to 17% of non-significantly variable genes (supplementary Fig.S10). The genes which are highly variable and differentially methylated are spread across all developmental comparisons (supplementary Fig.S10). A GO enrichment of these genes compared to all highly variable genes shows enrichment for various processes, including “*mRNA cis splicing, via spliceosome*” (GO:0045292), “*tissue remodeling*” (GO:0048771) and “*regulation of cell development*” (GO:0060284) (supplementary 1.0.7).

## Discussion

We tracked genome-wide DNA methylation dynamics throughout bumblebee development, from female worker somatic tissue and reproductive tissue into their male offspring larvae, pupae and adult head/brain tissue and then into the adult male sperm. We find that levels of DNA methylation vary dramatically in the reproductive tissue/gametes compared to the developing larvae/pupae head and adult brain. Ovaries show a sharp decrease in DNA methylation levels whereas sperm shows considerably higher DNA methylation levels. Interestingly, the high levels of DNA methylation in sperm is characterised by a more bimodal pattern of high/no DNA methylation in genes compared to intermediate levels. We also find differentially methylated genes in sequential developmental comparisons, most of which are lowly methylated genes and many are involved in processes relevant to each specific developmental stage / tissue. Finally, we find genes which have the most variable DNA methylation profiles throughout development are enriched for metabolic processes and also have a role in determining lifespan.

### Possible female-specific DNA methylation erasure

One of our key findings is that the major global differences in DNA methylation are seen between the somatic tissue across all stages and the reproductive tissue/gametes. DNA methylation is known to be tissue-specific in many species (Pai *et al*., 2011) and so this is unsurprising. However, we specifically find that ovaries show a marked decreased in overall DNA methylation, with sperm showing higher levels. Based on this finding, we hypothesise that DNA methylation profiles may be erased during oogenesis but remain stably transmitted through the male germline. Previous work in honeybees has shown no evidence for DNA methylation erasure during embryogenesis (Yagound *et al*., 2020).

However, Yagound *et al*. (2020) only examined DNA methylation inheritance from the point of view of transmission of the male’s methylome to his daughters. Transmission of the maternal methylome was unexamined. Cardoso-Júnior *et al*. (2021) do find similar DNA methylation profiles between the brains and ovaries of honeybee queens, contrary to what we find in bumblebees. Additionally, in stark contrast to our findings, Drewell *et al*. (2014) find both male and female honeybee gametes show higher methylation compared to male thorax tissue. This could suggest that if developmental erasure is occurring during oogenesis in bumblebees, it is not conserved between the two species. However, a controlled study using identical samples from both species would be needed to confirm maternal methylome persistence or erasure.

The high global levels of DNA methylation in sperm is characterised by higher numbers of genes showing high/no methylation rather than intermediate levels. Based on the above hypothesis of paternal methylome transmission, we suggest that this profile in sperm may represent a ‘pristine’ methylome. DNA methylation has been strongly linked to the underlying genotype in multiple hymenoptera species (Yagound *et al*., 2019; Wang *et al*., 2016), including bumblebees (Marshall *et al*., 2019). Additionally, Harris *et al*. (2019) also find evidence for a highly maintained sperm methylome, compared to somatic tissue in honeybees. They suggest the stronger signal of DNA methylation in sperm is present to maintain signatures across generations. We speculate that the sperm methylome may represent the genotype-mediated DNA methylation profile, which then later diversifies by tissue, developmental stage, age, environmental exposure etc. It may also be that we see more genes with low/medium levels of DNA methylation in head and brain tissue due to a loss of accurate transmission of epigenetic information through mitosis. This loss of epigenetic information over time is documented in other species and is strongly associated with ageing in mammalian systems (Yang *et al*., 2023). Sequencing of individual samples, with a known genetic background, would allow further testing of this hypothesis.

The idea of female-specific erasure based on our results is just one hypothesis for what we observe. During mammalian gametogenesis, there are waves of methylation and de-methylation in the gametes (Rotondo *et al*., 2021). It is possible we captured a specific stage of gamete development in males (only mature sperm). However, this is unlikely to be the case for the female ovary samples as they contain eggs of varying stages of development. If DNA methylation is indeed only transmitted through the paternal lineage in bumblebees, this would have implications for potential imprinting mechanisms (see Pegoraro *et al*. (2017) for a theoretical review of imprinting in bees) and for a possible role of heritable DNA methylation in environmental adaption (reviewed in Skinner and Nilsson (2021)).

### DNA methylation and bumblebee morphogenesis

In additional to global changes in DNA methylation between tissues and developmental stages, we have identified specific differentially methylated genes using sequential comparisons. Whilst these genes are involved in a multitude of processes, we find developmentally relevant differentially methylated genes throughout, which is suggestive of a functional role for DNA methylation in bumblebee development. To highlight one particular example, genes involved in larval development and imaginal discs are found to be differentially methylated between ovaries (containing developing eggs) and larvae. We find GO terms enriched for imaginal disc pattern formation, morphogenesis and imaginal disc-derived appendage development. Imaginal discs are tissue aggregations in holometabolous insects that develop into various adult structures during metamorphosis (Beira and Paro, 2016). Although we chose to apply a site and gene-level filter to consider a gene differentially methylated, perhaps also noteworthy may be genes excluded on that basis but with single differentially methylated loci. In the comparison between larvae and pupae such genes included *patched, smoothened, tramtrack* and *split ends*, which are linked with developmental processes such as compound eye morphogenesis, wing morphogenesis, nervous system development and Wg pathway regulation in larval tissues (supplementary 3.0.0), and the unfiltered results list showed GO enrichment for terms such as “*anterior/posterior lineage restriction, imaginal disc*” (GO:0048099) and “*Bolwig’s organ morphogenesis* (GO:0001746). Although the relationship between gene methylation and expression in invertebrates is uncertain (see below), it is well established in mammals that methylation changes in a single CpG is sufficient to impact expression (Zhang *et al*., 2010; Kitazawa and Kitazawa, 2007) and we do not rule out the possibility that these results may have biological significance. Overall, the identification of differentially methylated genes between developmental stages involved directly in the metamorphic process is suggestive of a functional role for DNA methylation in bumblebee development, though we cannot determine from our data whether it drives some regulatory process for these genes or if it is a consequence of gene regulatory changes occurring within development. Nevertheless, this finding allows for future work to examine the functional role of DNA methylation in development, for example by RNAi knockdown of DNMTs or though chemical de-methylation of the genome.

We also find the majority of differentially methylated genes between developmental stages show overall low levels of DNA methylation. Previous work has also found this is the case in DNA methylation differences between bumblebee castes (Marshall *et al*., 2023). This finding supports the idea that lowly methylated genes are able to promote phenotypic plasticity through creating more variable gene expression (Roberts and Gavery, 2012), whereas highly methylated genes are associated with housekeeping functions and stable gene expression across insects (Provataris *et al*., 2018). A recent hypothesis for the function of DNA methylation in terms of its response to environmental change has been proposed by Dixon *et al*. (2018). The ‘seesaw’ hypothesis suggests that transcriptional plasticity is brought about through lowly methylated genes increasing methylation levels as a group, and highly methylated genes decreasing methylation levels. This could explain why there are higher numbers of genes with intermediate levels of DNA methylation in the developing larvae and adult head tissue compared to the gametes, as this is where higher variability in gene expression is required for development.

Differential DNA methylation has generally not been linked to direct differential gene expression in invertebrates (Dixon and Matz, 2022). However, DNA methylation has been linked to a suite of other processes which may themselves govern downstream gene expression changes. For example, DNA methylation has been linked to alternative splicing in honeybees (Flores *et al*., 2012). Previous work has established large numbers of differential alternative splicing events throughout bumblebee development (Price *et al*., 2018). Some of the genes with the most variable DNA methylation in our study are indeed involved in mRNA splicing. DNA methylation changes may therefore be playing some role in mediating developmental plasticity through splicing. Additionally, other epigenetic mechanisms have been associated with insect metamorphosis, including microRNAs (Burggren, 2017) and histone modifications. Specifically, histone acetylation has been implicated in the metamorphic process in some fruit fly and butterfly species (Carré *et al*., 2005; Zhang *et al*., 2019; Mukherjee *et al*., 2012). DNA methylation in the silk moth influences histone acetylation which then impacts gene expression (Xu *et al*., 2021). This highlights the complex interplay between multiple mechanisms. A multi-omics study carried out on the same samples (to account for the effects of genetic variation) is required to disentangle the relative roles of epigenetic processes in mediating bumblebee metamorphosis.

Recent advancements in technology would allow a more precise look at how DNA methylation may be involved, either directly or indirectly, in mediating bumblebee development. For example, single-cell methylation sequencing would allow cell-specific profiles to be determined. Indeed, a recent paper has developed a method to obtain single-cell methylomes from mouse brain tissue (Liu *et al*., 2023). CUT&Tag methods allow the identification of histone modifications using considerably lower DNA input (Kaya-Okur *et al*., 2019), making this technology more accessible to small insect tissue samples. Finally, in order to determine if there is a causative role for DNA methylation in bumblebee development, experimental knockouts using RNA-i or specific DNA methylation additional/removal using CRISPR-Cas9 (Vojta *et al*., 2016) are required. This study lays the groundwork required for these future research avenues.

## Acknowledgements

BJH and MP were funded by NERC grant NE/N010019/1, awarded to EBM. HM was funded by the Leverhulme Trust Research Project RPG-2020-363, awarded to EBM. This research used the ALICE2 High Performance Computing Facility at the University of Leicester.

## Data accessibility

All sequencing data related to this project can be found under NCBI BioProject PRJNA573598. Custom scripts for the genome-wide analysis can be found at https://github.com/MooHoll/bumblebee_developmental_meth.

## Authors’ contributions

EBM and MP designed the study. MP carried out husbandry, sampling and DNA extraction. BJH and HM analysed the data. BJH and HM wrote the initial draft. All authors were involved in redrafting.

