## Supplementary_2 for "A role for DNA methylation in bumblebee morphogenesis hints at female-specific developmental erasure"

### Supplementary figures

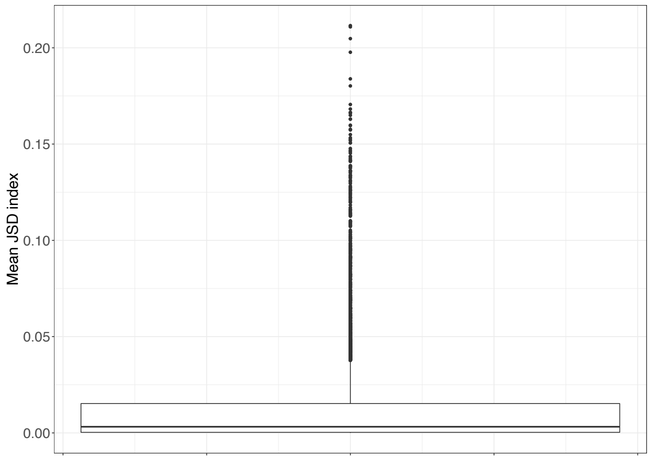

Figure S1: Distribution of Jensen-Shannon diversity (JSD) indices for all genes tested (n = 9,016). Genes were classes as showing a significant JSD index if they were greater than 1.5x the interquartile range, i.e. an outlier on this plot.

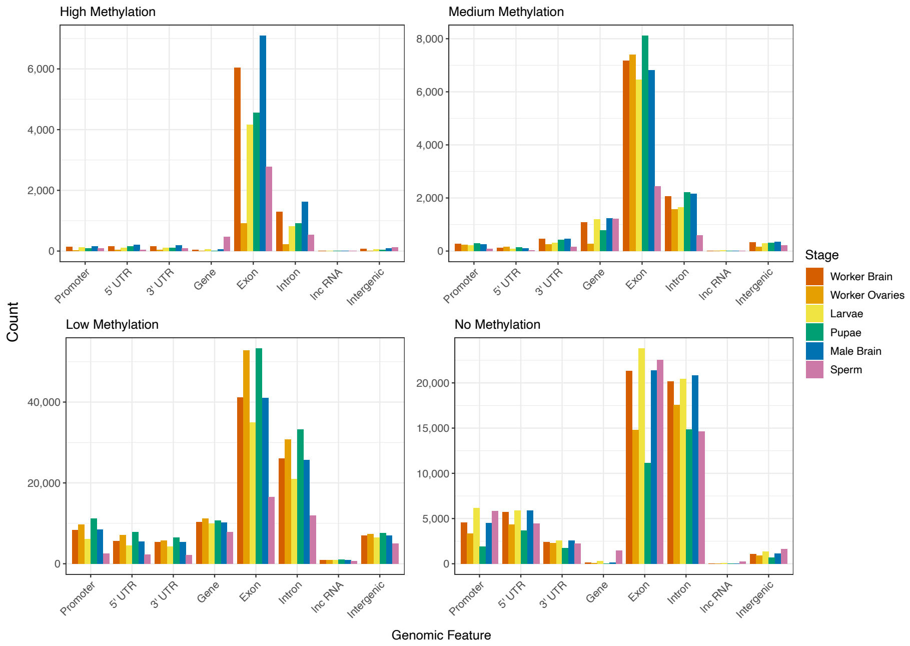

(a)

Figure S2: Number of features per developmental stage that are classed as showing high methylation (>0.7), medium methylation (>0.3-0.7), low methylation (>0-0.3) and no methylation.

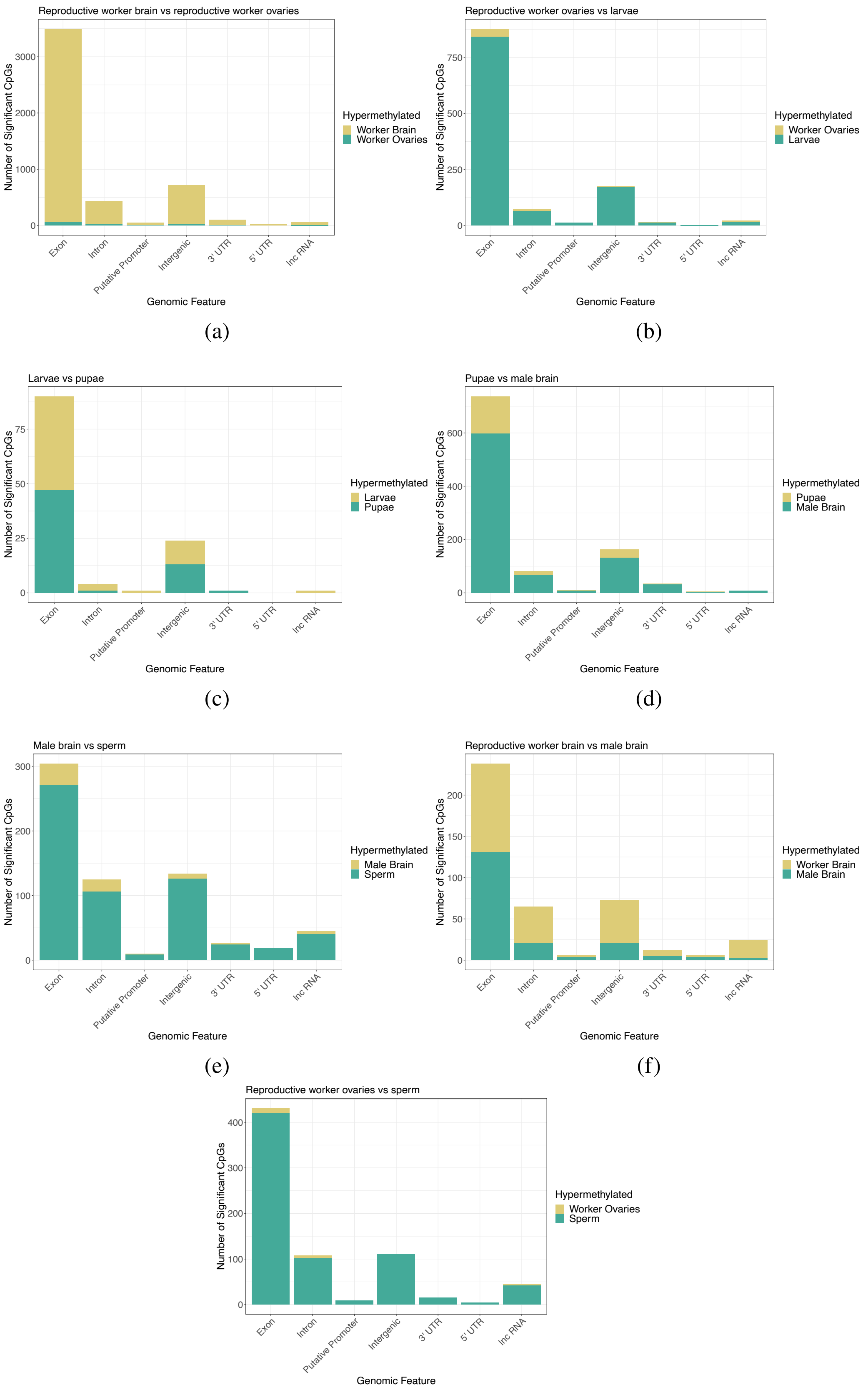

Figure S3: The genomic location of the differentially methylated CpGs identified in each comparison coloured by the hypermethylated developmental stage.

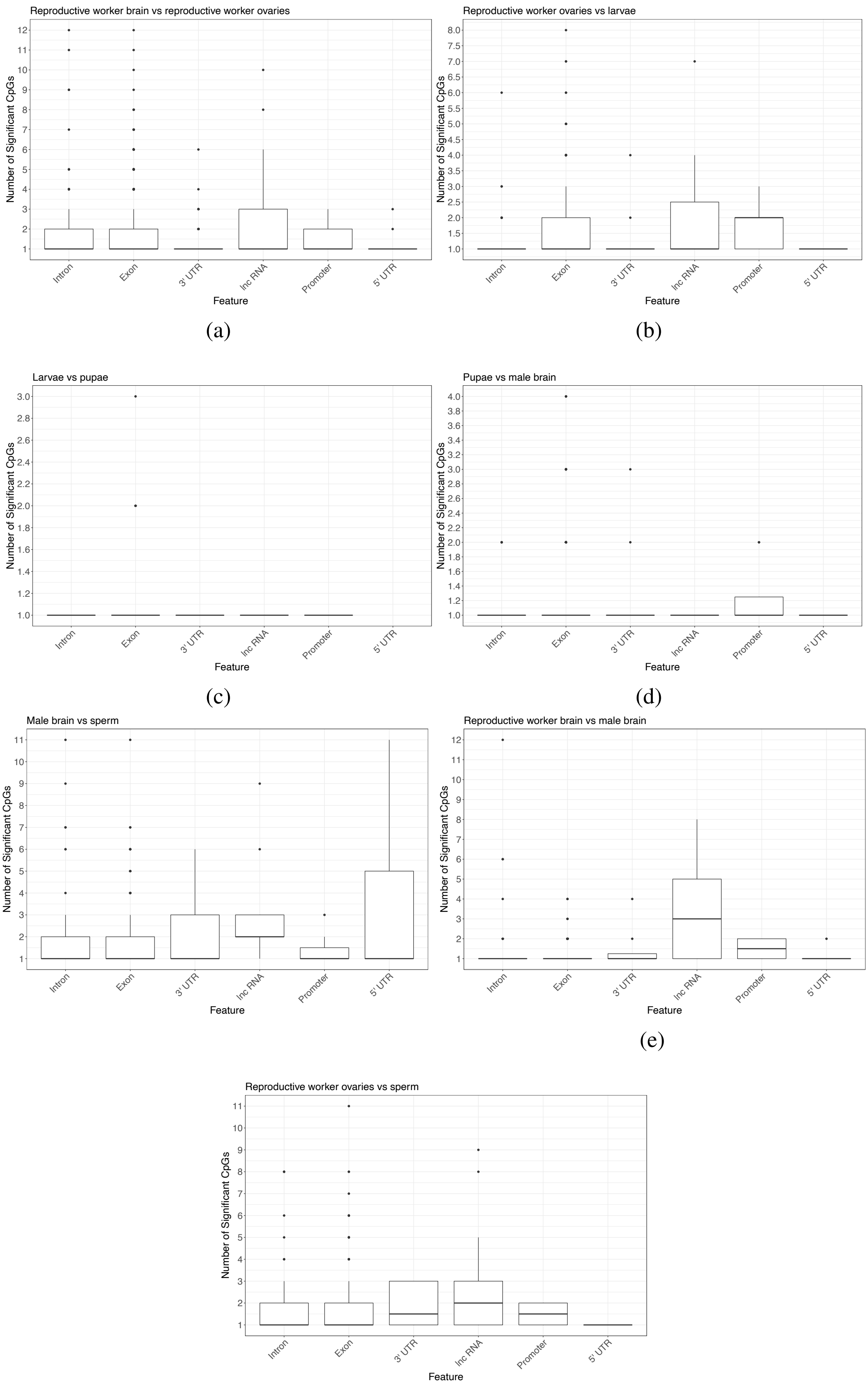

Figure S4: Number of differentially methylated CpGs found within each genomic feature per developmental stage comparison.

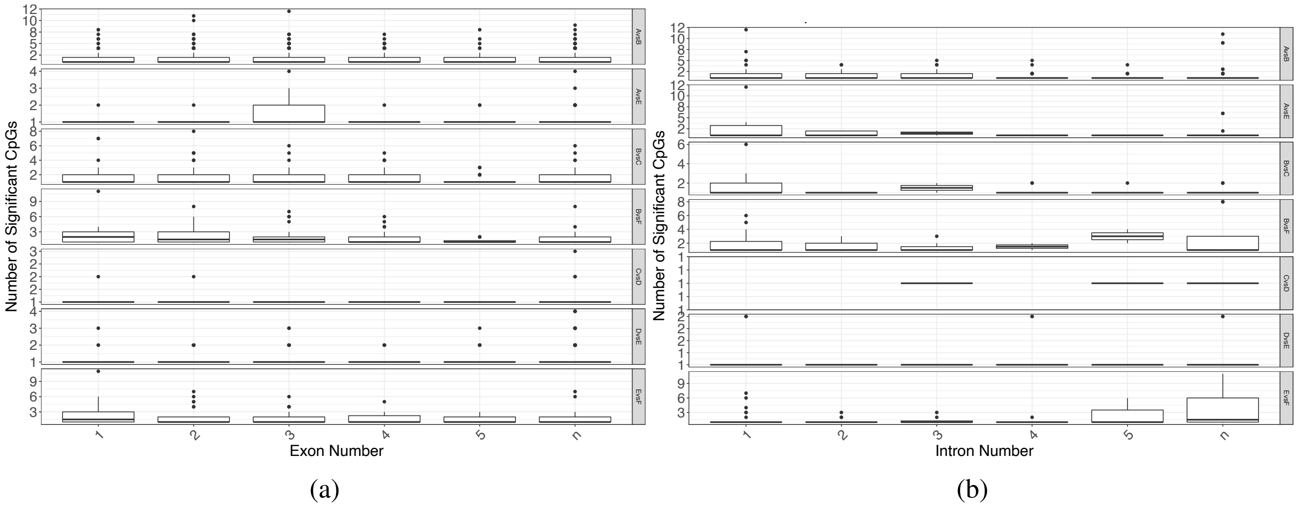

Figure S5: Number of differentially methylated CpGs found within the first five exons (a) and introns (b), with exons and introns 6+ represented by n. The letters represent each developmental stage, A = reproductive worker brain, B = reproductive ovaries, C = larvae, D = pupae, E = male brain and F = sperm.

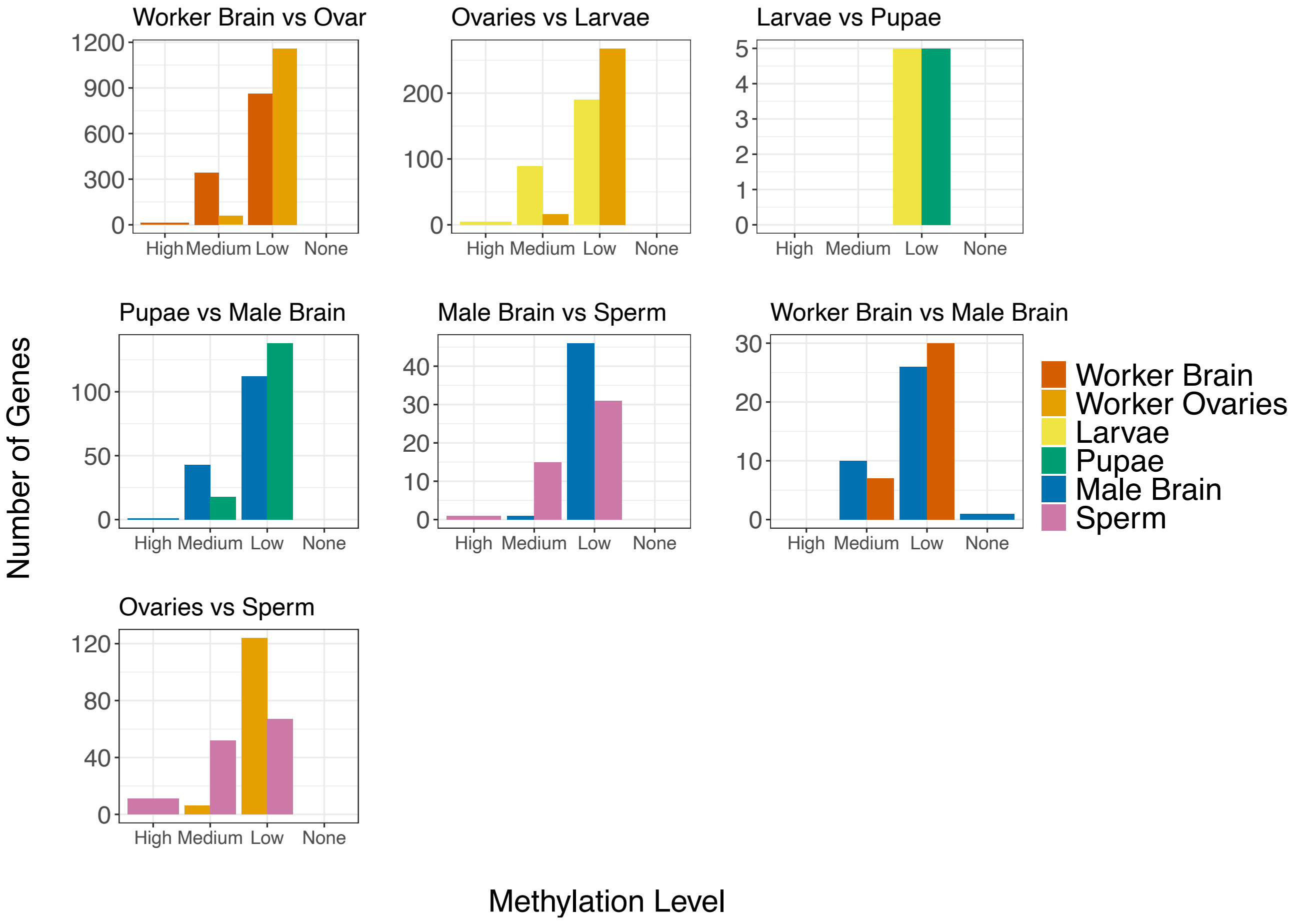

(a)

Figure S6: Differential methylation comparison showing the weighted methylation level of the differentially methylated genes. High >= 0.7, medium < 0.7 and >=0.3, low < 0.3 and > 0, none is 0.

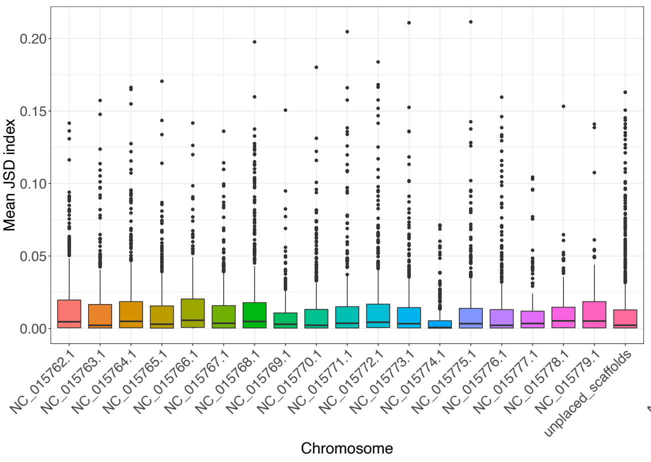

(a)

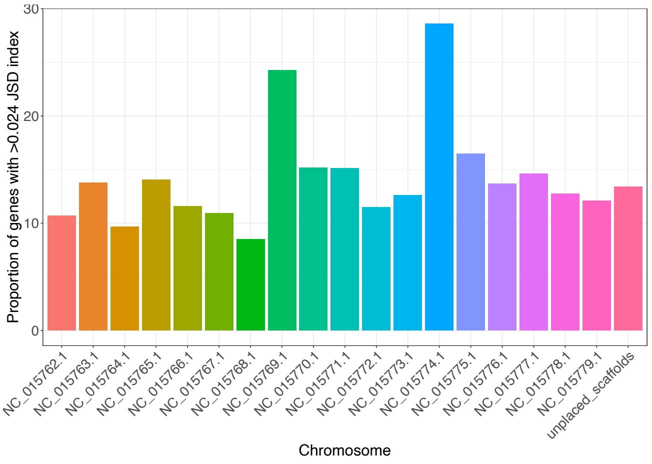

(b)

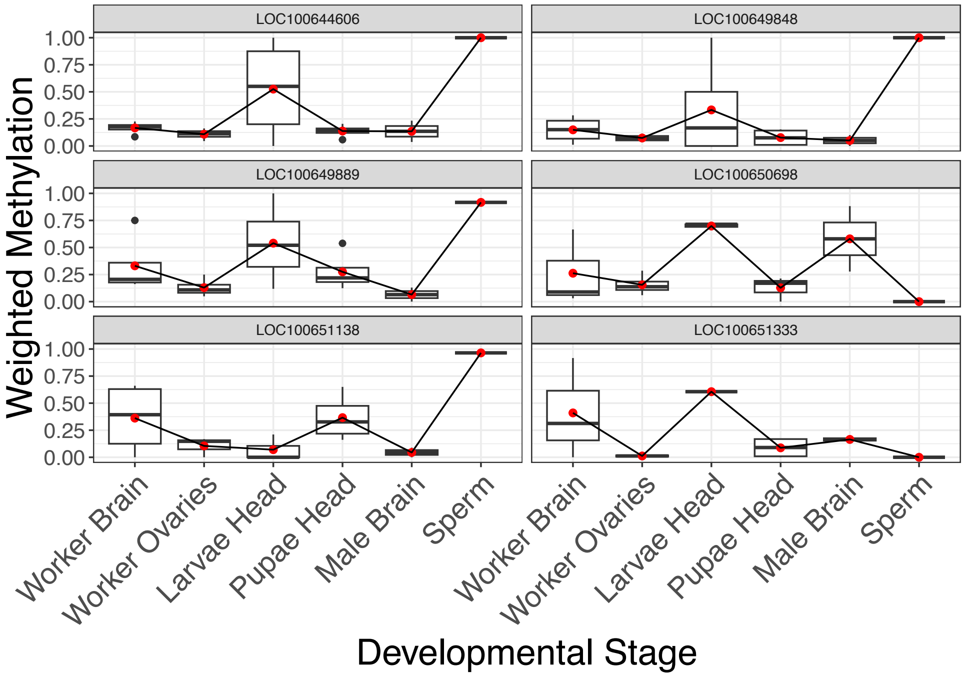

(c)

Figure S7: (a) Jensen-Shannon diversity indices for every gene, plotted by chromosome. (b) Proportion of genes per chromosome which show a significant Jensen-Shannon diversity index, i.e. have more highly variable DNA methylation states across developmental states compared to most other methylated genes. (c) Weighted methylation level of the top six most variable genes. The red dot shows the mean across replicates within each developmental stage.

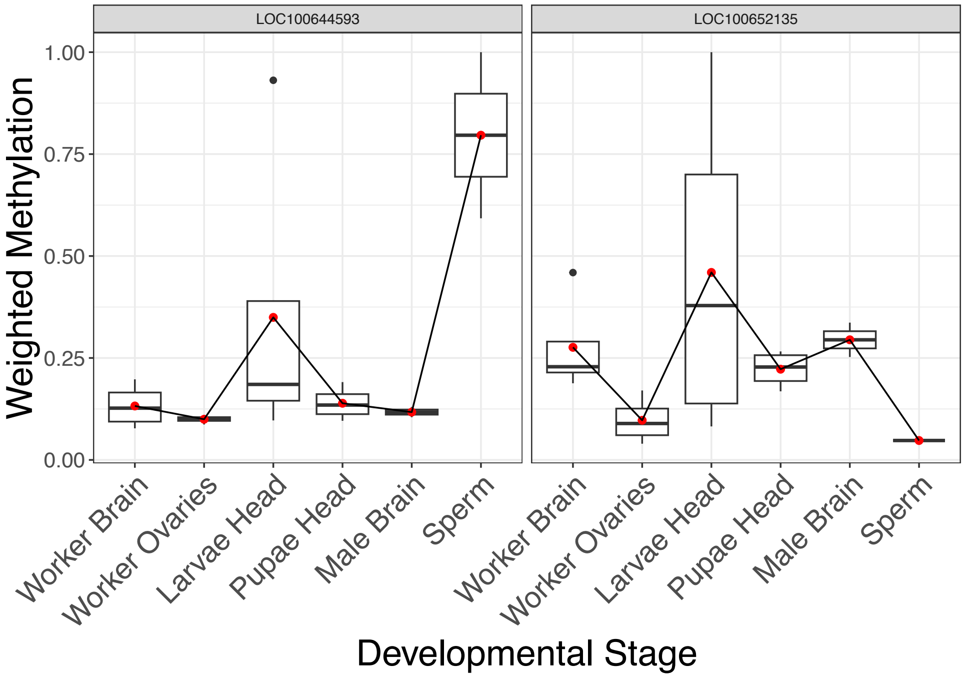

Figure S8: Weighted methylation level of two genes involved in life span. The red dot shows the mean across replicates within each developmental stage. LOC100644593: ras-like protein 1. LOC100652135: caspase-6.

Table S1: The number of hypermethylated CpGs per developmental stage for all comparisons. A chi-squared goodness of fit was carried out per comparison to determine if there are significantly more hypermethylated CpGs in oen developmental stage compared to the other.

| | **Worker Brain** | **Worker Ovaries** | **Larvae Head** | **Pupae Head** | **Male Brain** | **Sperm** | **X-squared** | **df** | **p** |
|---|---|---|---|---|---|---|---|---|---|
| **Worker Brain vs Worker Ovaries** | 3819 | 91 | | | | | 3554.5 | 1 | <0.001* |
| **Worker Ovaries vs Larvae Head** | | 47 | 945 | | | | 812.91 | 1 | <0.001* |
| **Larvae Head vs Pupae Head** | | | 51 | 48 | | | 0.0909 | 1 | 0.763 |
| **Pupae Head vs Male Brain** | | | | 147 | 654 | | 320.91 | 1 | <0.001* |
| **Male Brain vs Sperm** | | | | | 52 | 575 | 436.25 | 1 | <0.001* |
| **Worker Brain vs Male Brain** | 163 | | | | 150 | | 0.5399 | 1 | 0.462 |
| **Worker Ovaries vs Sperm** | | 20 | | | | 381 | 623.28 | 1 | <0.001* |

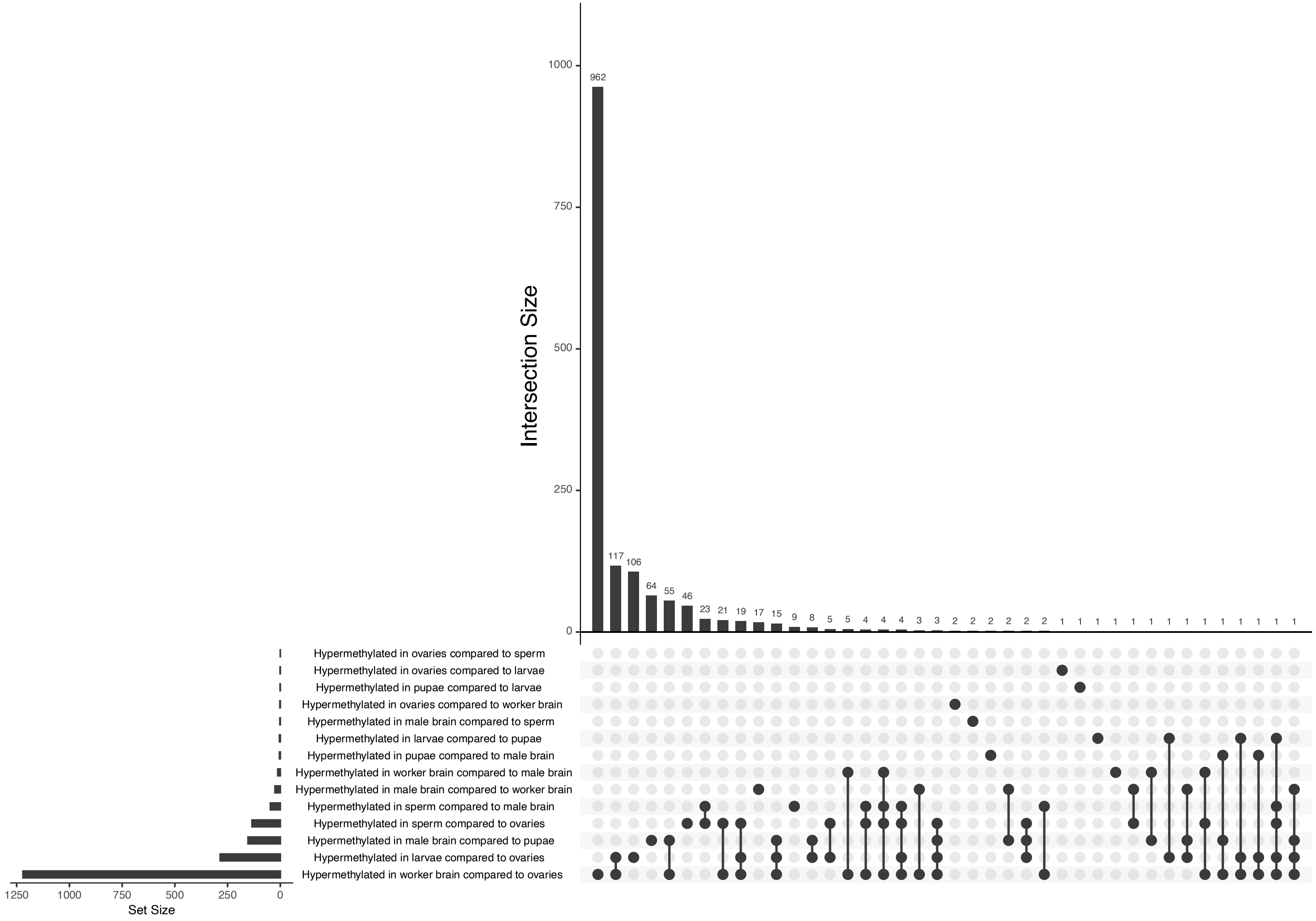

Figure S9: Overlap of genes between differential DNA methylation comparisons. The set size indicates the total number of hypermethylated genes, the intersection size shows how many of those are common between sets, as indicated by the connections in the bottom panel. E.g. 117 genes hypermethylated in worker brains and larvae compared to ovaries.

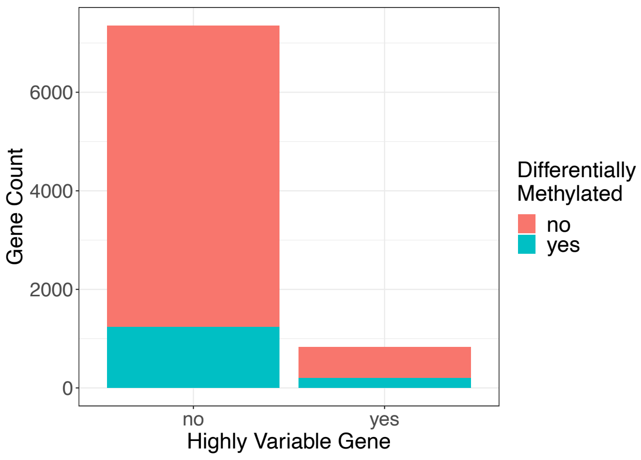

(a)

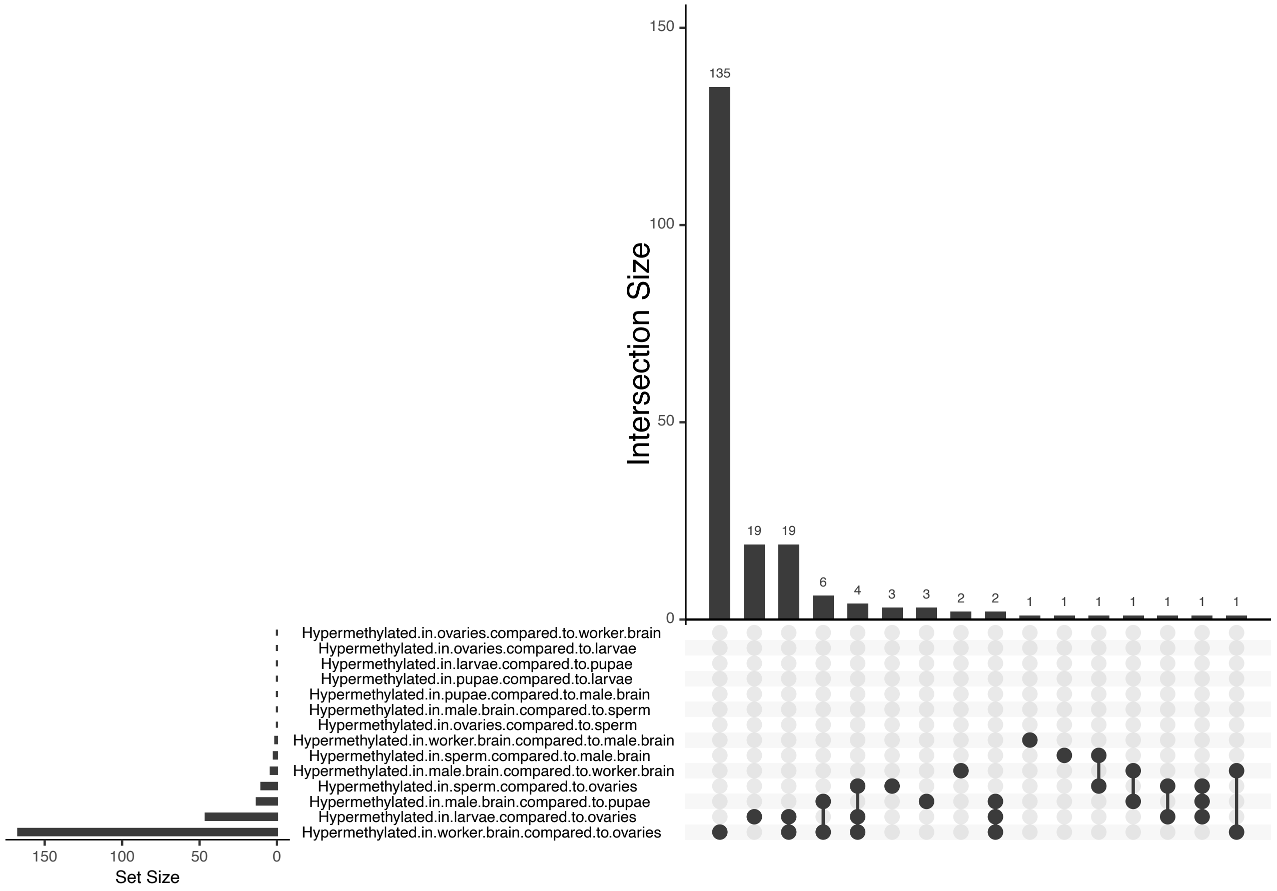

(b)

Figure S10: (a) Number of genes which are differentially methylated in any comparison and also show high variability across developmental stages. (b) UpSet plot showing in which comparison the number of genes which are both differentially methylated and also highly variable are found.
